# Lessons learned from comparing molecular dynamics engines on the SAMPL5 dataset

**DOI:** 10.1101/077248

**Authors:** Michael R. Shirts, Christoph Klein, Jason M. Swails, Jian Yin, Michael K. Gilson, David L. Mobley, David A. Case, Ellen D. Zhong

## Abstract

We describe our efforts to prepare common starting structures and models for the SAMPL5 blind prediction challenge. We generated the starting input files and single configuration potential energies for the host-guest in the SAMPL5 blind prediction challenge for the GROMACS, AMBER, LAMMPS, DESMOND and CHARMM molecular simulation programs. All conversions were fully automated from the originally prepared AMBER input files using a combination of the ParmEd and InterMol conversion programs.

We find that the energy calculations for all molecular dynamics engines for this molecular set agree to a better than 0.1% relative absolute energy for all energy components, and in most cases an order of magnitude better, when reasonable choices are made for different cutoff parameters. However, there are some surprising sources of statistically significant differences. Most importantly, different choices of Coulomb’s constant between programs are one of the largest sources of discrepancies in energies. We discuss the measures required to get good agreement in the energies for equivalent starting configurations between the simulation programs, and the energy differences that occur when simulations are run with program-specific default simulation parameter values. Finally, we discuss what was required to automate this conversion and comparison.

## INTRODUCTION

The goal of the ongoing SAMPL blind prediction challenges [1–4] is to compare purely computational blind predictions of thermodynamic properties, such as hydration free energies, partition coefficients, and binding free energies, for a range of both model and more realistic systems. Such blind prediction challenges can be very useful in identifying unexpected reasons for differences between methods that should, in theory, yield the same result. For example, the same program used with what is listed as the same force field can sometimes still yield significantly different results. In the SAMPL4 blind test, two different sets of simulations performed with GROMACS, TIP3P water, and GAFF/AM1-BCC parameters had differences of 2 *±* 0.5 kJ/mol that were ultimately tracked down to whether the large host molecule had AM1/BCC partial charges determined fragment-wise or for the entire molecule at the same time. This level of detail often does not make it into publication, [5] which can severely hamper efforts in reproducing results. During the SAMPL5 challenge reported in this special issue, free energies of binding using the same GAFF/RESP force field with TIP3P water were calculated with a number of different molecular dynamics engines, often with statistically signficant differences, but it is still not entirely clear where those differences come from [6].

One particular question that can be difficult to address is to what extent methods that are supposed to be identical will give different results when attempted with different simulation programs. Therefore, one of the tasks carried out in preparation for SAMPL5 was to prepare starting simulations in several of the most common molecular simulation packages (AMBER [7], GROMACS [8], LAMMPS [9], and DESMOND [10]). To ensure that the simulation inputs were translated correctly between programs, it was also necessary to compare the evaluated energies of the initial configurations in the simulation programs that were native to each file format to ensure that the translation had been done correctly. This also involved determining the necessary simulation conditions and parameter choices for each program to give the same, or sufficiently the same, energy. To make this task feasible for the 22 host-guest systems and 190 distribution coefficient systems, this process was necessarily highly automated. We present here the results of energy comparisons carried out in the process of SAMPL5 preparation and validation. One important change from the initial work carried out for SAMPL5 and this paper is adding CHARMM-format starting files as well as the initial energies generated with the CHARMM simulation program.

There are two main ways that one can compare molecular simulation engines. The first task that a molecular simulation engine has is to take a molecular configuration and a model (i.e. a specification of all force field parameters) and from these ingredients generate the energy of the model in that configuration and, in molecular dynamics approaches, also the forces acting on each particle. Next, given the assignment of energies and forces to a configuration, a molecular simulation engine then also generates a sequence of configurations that belong to a desired ensemble of that model, such as the microcanonical (NVE), canonical (NVT), or isobaric-isothermal (NPT) ensemble, with their corresponding probability distribution for each configuration. There are therefore two types of comparison that can be done between simulation engines. The first is to take one (or a few) configurations and compare the potential energies engines generate. The second is to compare the observables, such as density or enthalpy, that the simulations generate after the generation of an ensemble of configurations. The comparison of the final experimental observables calculated from simulation ensembles is the most fundamentally important type of comparison, since it corresponds directly to experiment.

However, this second task requires a large number of different decisions that are made mostly independently of the assignment of energy to a set of coordinates. For example, slightly different integration methods will give rise to slightly different ensembles. Different thermostats will converge to the correct ensemble (if they are actually correctly implemented thermostats) but the speed at which they approach to that value can vary. Different programs have different recommended integration schemes.

Additionally, simulation observables are ensemble averages, and are thus statistical quantities with associated statistical error. Since the uncertainty scales as (simulation time)^−1*/*2^, increasing the accuracy by a factor of 10 requires 100 times as much simulation time. Small differences in the parameters used to run the simulations give rise to similarly small changes in the ensemble averages. If we attempt to calculate a small difference between ensemble averages 〈*O*〉_1_ and 〈*O*〉_2_, carried out at simulation parameter sets 1 and 2, with statistical uncertainties *σ*_1_ and *σ*_2_, the error in 〈*O*〉_1_ − 〈*O*〉_2_ will be 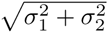. If 〈*O*〉 is, for example, the enthalpy of a calculation, it might be of order −100, 000 kJ/mol for a large simulation. For a given amount of simulation time, if the relative error in 〈*O*〉 is 0.0001% or about 10 kJ/mol, then error in 〈*O*〉_1_ − 〈*O*〉_2_ will be of order 14 kJ/mol. Clearly, it would be important to know if a change in a simulation parameter changed the enthalpy difference by anywhere near 14 kJ/mol. To take that uncertainty down to, say, a 95% confidence interval of 1 kJ/mol would take approximately (2 standard deviations *×* 14)^2^ ≈ 800 times as much simulation as determining 〈*O*〉 itself to one part in 10^−6^. Reweighting approaches have recently been developed to include the correlation between variables, allowing in many cases the uncertainty to be calculated by one to three orders of magnitude more efficiently. [11] However, even with this acceleration, it is still extremely expensive.

Because of the difficulties of comparing the simulation ensemble averages, as outlined above, this study focuses on validation of the first task of molecular simulation engines, the generation of energies from a configuration of a molecular system and a well-defined model for that molecular system, testing only those parts of the molecular dynamics simulation engines. Addressing the multitude of possible ways that simulations could differ using different integration schemes and running simulations long enough to detect small changes is beyond the scope of this study. The validation of the energy generation presented here serves as a necessary building block for later studies to more easily evaluate the differences between simulation engines in calculating simulation observables, and the comparison of more advanced simulation methods between different simulation programs.

Comparison between simulation programs are typically tedious and error prone, because the input configurations and model must be converted. This is either done using painstaking manual copy-and-pasting, one-off scripts, or occasionally existing scripts that can convert from one specific program to another. Some examples include ACPYPE, a converter from AMBER to GROMACS [12], CHAMBER [13] a converter from CHARMM to AMBER, TopoGromacs, a converter from CHARMM to GROMACS [14], CHARMM-GUI [15] which can convert from CHARMM to GROMACS (and uses ParmEd to manipulate AMBER files), and amber2lmp, a script converting between LAMMPS and AMBER files. However, converting between a large number of simulation programs has not, to the best of our knowledge, previously been done in a study.

A significant additional challenge in comparing the output of programs with theoretically the same model occurs when the energy and forces are approximated in order to increase the number of molecular dynamics or Monte Carlo steps in a fixed amount of computing time. For example, Lennard-Jones terms may be truncated at some separation distance distance with some sort of approximation for longer distances [16–18], or Coulombic interactions long-range terms may be approximated by an interpolated mesh [19] rather than a direct lattice sum. Not only does each program make different default choices, most of these choices are left up to the user, meaning different results can be obtained by different users of the same code, and the recommended or default behavior of each code will almost certainly differ from program to program to some degree.

This study therefore focuses on the automated conversion of molecular simulation input files using (to the extent possible) automated all-to-all conversion tools, and the comparison and validation of the energies of single configurations among these programs. In this process, we attempt to find reasonable simulation parameter choices that allow the nonbonded energies to be directly compared.

## METHODS

The molecular interconversion software programs InterMol (https://github.com/shirtsgroup/InterMol) and ParmEd (http://github.com/ParmEd/ParmEd) were used to perform comparisons between five different simulation input parameter files and engines. InterMol is designed as a generalizable all-to-all converter between molecular simulation file formats; however, it currently only has full support for GROMACS, LAMMPS, and DESMOND file formats. ParmEd is a library for defining and manipulating atomic-level molecular topologies with force field descriptions. It provides a program-agnostic representation of a molecular topology and its force field that supports editing molecular topologies as well as providing the infrastructure to convert files between the native formats for the GROMACS, AMBER, CHARMM, and OpenMM programs.

We took advantage of this overlap in conversion functionality to provide output files in five formats. The process is as follows: We took files initially parameterized in AMBER format with GAFF/RESP force field parameters using AmberTools, as described in the SAMPL5 overview paper [6] and read them using ParmEd. We then used ParmEd to convert them into GROMACS input file formats. We then convert from these GROMACS files into LAMMPS and DESMOND input files using InterMol. ParmEd was also used for this study (though not the original SAMPL5 release) to convert the AMBER simulation files into CHARMM simulation files directly.

We use the InterMol `convert.py` tool to manage all of the conversions (including interfacing with the ParmEd API). InterMol allows control of simulation input parameters by either reading a user-defined (or default) sample simulation parameter file (for DESMOND, AMBER, and GROMACS) or inserting user-defined strings defining nonbonded terms into the parameter and topology files (LAMMPS and CHARMM). Full any-to-any conversion is not yet possible using the combination of tools so far, since ParmEd cannot yet convert between some dihedral formats, making it impossible to write many valid GROMACS files into CHARMM or AMBER formats.

We use the 22 host-guest molecules distributed as part of the SAMPL5 blind challenge for our comparison of energies. The systems used for the distribution coefficient challenge portion of SAMPL5 were also converted and their energies evaluated [20], but the results were similar with no additional lessons, and so we focus on the host-guest molecules here. Both energies and configurations for the distribution coefficient challenge are for the near future still posted on the SAMPL5 web site (https://drugdesigndata.org/about/sampl5).

The first two hosts, with six guests apiece, are OAH [21] and OAMe [22, 23] from the Gibb laboratory, are also known as octa-acid (OA) and tetra-endo-methyl octa-acid (TEMOA). The last host is CBClip [24], with 10 ligands, from the Isaacs laboratory. Specific details of the topology construction and configuration generation is described in the SAMPL5 overview paper. [6]

For this study, we attempted to use the most up-to-date releases of all molecular dynamics simulation programs. In most cases, this resulted in very little difference in the results between the current study and the SAMPL5 study, but in some cases as noted, the results do change. Units are given in SI units (kJ/mol and nm, for example), though different programs use different default units.

The five programs were:

- AMBER: Energies were calculated originally for SAMPL5, with sander as included AmberTools 14, but for the current study sander from the most recent AmberTools 16 were used.
- GROMACS: Energies were calculated originally and here with GROMACS 5.0.4, compiled in double precision.
- DESMOND: Energies were calculated in the original SAMPL5 release with version 3.6012 (distributed as part of the Schrödinger 2013 package for academic use) but all tests are performed here with version 4.5 (distributed as part of the Schrödinger 2016-1 package for academic use). Rather than directly writing the DESMOND .dms files, the automated conversion routines were written to create MAESTRO .cms files.
- LAMMPS: Energies were calculated with the April 5, 2014 build in the SAMPL5 release, and the Feb 16, 2016 release in the current study. Only the additional modules to run atomistic simulations were installed (`MOLECULE`, `RIGID`, and `KSPACE`)
- CHARMM: Energies were only generated for this study, with developmental version 40b2 of the free version of CHARMM (“charmm”), which has all of the features of the CHARMM program except for the DOMDEC and GPU high performance modules.

All programs were compiled in RHEL 7 with the gcc 4.8.5 compiler suite, and run on the same desktop computer.

It is difficult to choose simulation parameters that agree among all simulation engines. For example, each program generally has different types of default switching functions to taper nonbonded interactions. There is certainly an argument to be made that any parameters that affect the energy, such as switching scheme, cutoff, and Coulomb’s constant should be considered part of the forcefield, but in essentially all common usage, they are not—‘force field’ in general usage refers to the specification of partial charges, atom types, van der Waals parameters, mixing rules, and bond/angle/torsion parameters.

For this comparison, we tried to choose for our nonbonded interactions methods that were sufficiently cutoff independent that differences in the cutoff scheme between programs would minimally affect the results, given the limitations imposed by the size of the system.

For electrostatic interactions, we chose either particle mesh Ewald (PME) implementations (CHARMM, AMBER, GROMACS, CHARMM) or particle-particle particle-mesh (PPPM) methods (LAMMPS). We chose a cutoff of 1.5 nm for both Coulomb and van der Waals interactions to eliminate much of the issues with errors at short range cutoffs. We chose a real space error cutoff of 1 *×* 10^−8^, which corresponds to a *κ* (also known as *β*) parameter of 0.020822755 nm. For PPPM, we chose a tolerance of 1 *×* 10^−8^. For AMBER and CHARMM, we used a PME mesh grid of 48 *×* 48 *×* 48 grid points with 4th order interpolation. DESMOND allows significantly less control over the PME parameters at the MAESTRO interface level, and we used an PME relative tolerance (`ewald_tol`) of 1 *×* 10^−10^.

For Lennard-Jones interactions, we avoid the problems of trying to match switching schemes between programs, which are usually quite different, by using an abrupt cutoff to zero potential. This approach is not recommended for running molecular dynamics simulations, as it creates a mismatch between forces and energies, but which is reasonable for comparing simulation energies. An analytic isotropic long-range correction was used for LAMMPS, AMBER, DESMOND, and GROMACS [16, 17], with the isotropic periodic sum approach [18] used for CHARMM. At this longer range, the results become essentially independent of the precise cutoff for both methods, though the isotropic periodic sum is less cutoff dependent, as seen in Table I. This cutoff independence in the Lennard-Jones energies is expected for systems that are homogeneous at long range, such as a host-guest system surrounded by water. However, this approach will not be cutoff-independent for a heterogeneous system such as a lipid bilayer or a liquid/vapor interface [25]. The parameters listed above are referred to in this study as the ‘ideal’ parameters, and listed in Table II.

**TABLE I:** Using an analytical correction, the van der Waals energy due to the Lennard-Jones interactions are essentially independent of cutoff, with a total change of 0.001% in the total van der Waals energy for the analytic long range correction in GROMACS (and similar to other programs) and 0.00008% with the isotropic periodic sum in CHARMM, over a change of 0.3 nm cutoffs. This particular example uses the CBC-G1 system.

| Distance (nm) | van der Waals Energy (kJ/mol) |  |
| --- | --- | --- |
|  | GROMACS | CHARMM |
| 1.5 | 12223.489 | 12223.753 |
| 1.4 | 12223.524 | 12223.749 |
| 1.3 | 12223.667 | 12223.748 |
| 1.2 | 12223.659 | 12223.742 |

**TABLE II:**
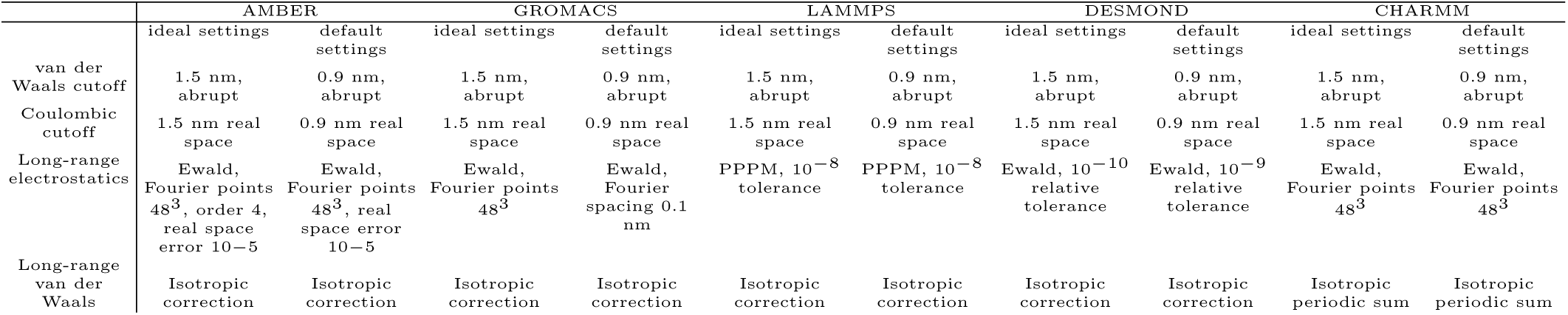
Key nonbonded parameters used in this study for both default and ideal energy validation test. *n*^3^ is shorthand of lattice numbers *n × n × n* in the *x*, *y* and *z* direction.

|  | AMBER |  | GROMACS |  | LAMMPS |  | DESMOND |  | CHARMM |  |
| --- | --- | --- | --- | --- | --- | --- | --- | --- | --- | --- |
|  | ideal settings | default settings | ideal settings | default settings | ideal settings | default settings | ideal settings | default settings | ideal settings | default settings |
| van der Waals cutoff | 1.5 nm, abrupt | 0.9 nm, abrupt | 1.5 nm, abrupt | 0.9 nm, abrupt | 1.5 nm, abrupt | 0.9 nm, abrupt | 1.5 nm, abrupt | 0.9 nm, abrupt | 1.5 nm, abrupt | 0.9 nm, abrupt |
| Coulombic cutoff | 1.5 nm real space | 0.9 nm real space | 1.5 nm real space | 0.9 nm real space | 1.5 nm real space | 0.9 nm real space | 1.5 nm real space | 0.9 nm real space | 1.5 nm real space | 0.9 nm real space |
| Long-range electrostatics | Ewald, Fourier points $48^3$, order 4, real space error 10–5 | Ewald, Fourier points $48^3$, real space error 10–5 | Ewald, Fourier points $48^3$ | Ewald, Fourier spacing 0.1 nm | PPPM, $10^{-8}$ tolerance | PPPM, $10^{-8}$ tolerance | Ewald, $10^{-10}$ relative tolerance | Ewald, $10^{-9}$ relative tolerance | Ewald, Fourier points $48^3$ | Ewald, Fourier points $48^3$ |
| Long-range van der Waals | Isotropic correction | Isotropic correction | Isotropic correction | Isotropic correction | Isotropic correction | Isotropic correction | Isotropic correction | Isotropic correction | Isotropic periodic sum | Isotropic periodic sum |

We also ran the test with an attempt to be as close to default parameters as possible and still have comparable energies. For all simulations, we used a 0.9 nm abrupt cutoff, as differences in switching methods lead to significantly different Lennard-Jones energies. For AMBER, we used the same Ewald parameters (48*×*48*×*48 Fourier points in the grid), with the isotropic analytical dispersion correction included for Lennard-Jones. For GROMACS, a Fourier spacing of 0.1 nm is used. For LAMMPS, we set pair style lj/cut/coul/long 9.0 9.0. We include a dispersion correction, and use PPPM, with 10^−8^ as the tolerance. For DESMOND, default ewald parameters were used, with ewald tol equal to 10^−9^. These parameters are referred to as the ‘default’ parameters in this study, and listed in Table II.

All input files used are documented in the “runfiles.tgz” directory of the Supporting Information (with details given in the README.txt). Converted input simulation files are available in AMBER, CHARMM, DESMOND, GROMACS, and LAMMPS formats for the 22 host-guest systems are provided as tar/gzipped files. The 20 configurations used in the paper for the comparison of average simulation energies are only included with the original AMBER `.rst7` files for storage space reasons.

We also examined how dependent the energy differences are on the individual configurations. For twelve of the systems (the six OAMe and the six OAH octa-acid host-guest systems), we take 20 different configurations. These configurations were generated with electrostatic parameters and Lennard-Jones parameters as listed in the ‘default’ parameter section. Temperature was maintained at 298.15 K with the Langevin thermostat with a damping constant *γ* of 1 ps^−1^ with a timestep of 2 fs with constrained bonds involving hydrogen. Simulations were started from the SAMPL5 example files and run for 2 ns, and configurations were taken every 100 ps.

Different simulation packages use file formats which have different default precisions, which means both input parameters and coordinates can be rounded differently. This truncation can have dramatic impacts on computed energies. Clearly, all parameters must match, or else the model will be different than intended. In InterMol, we use by default 8 decimal places in the parameters to ensure matches to high precision, though PARMED truncates CHARMM exported files at a varying precision depending on the parameter (almost always 5 significant digits, except for angle force constants, which were 4 significant digits). This did not cause adverse deviation for bond or angle energy terms compared to other programs, though may account for CHARMM dihedral energies being 3-4 times further from the program average than other programs. Although nonbondeds were truncated at 7 significant figures, no rounding with respect to the other programs occurred for this dataset.

However, matching the precision in the coordinates in two file formats is also important in order to verify energies. We examined the importance of matching the precision in coordinates. While keeping the precision of the input files the same, we truncated the precision of the converted files to a range of different precisions, ranging from four decimal places to nine decimal places to see how the precision of the energy components are affected.

Note that using lower precision coordinate files will not have an effect on ensemble averages if such files are only used as starting points for simulation. But the precision in the coordinates will matter quite a bit if stored output configurations are used to re-evaluate the energy contributions, as low precision coordinates could introduce significant error.

Another source of differences in energy relates to the precision of the binary. We therefore also compare the deviation from the program average for the same version of GROMACS compiled in double precision (the precision that is used in the reference calculations) and in a reduced precision version of the binary. In GROMACS, “mixed precision” is the official name for this reduced precision version, which uses single precision for the state vectors (velocity, position) and forces, but not for particularly sensitive calculations like the virial and the integration. It is essentially, therefore, single precision, and we will refer to this in the text, since the energies are being computed in single precision.

## RESULTS

We first compare ten different energy terms between the five different simulation programs with the previously defined ideal nonbonded calculation settings. Results are shown in Fig. 1. All results are averaged over the 22 host-guest systems included in the SAMPL5 blind prediction challenge. To avoid picking a favored reference program, we look at the deviation of each term from the average of all five programs, calling this the “program average” for the molecule. “Potential energy” is the total potential energy of the system, but is not the total over all 9 other listed energy terms, since several of these terms are sums of other terms: “Bonded” is the sum of “Bonds”, “Angles”, “All dihedrals” (including both improper and proper dihedrals). It is difficult to assign a label “proper” or “improper” to a given dihedral, since in different programs the same functional form is used for both types. We thus put all of these energies together as dihedral energy when reporting them to avoid having to deal with the ambiguity of different decompositions. “Electrostatic” is the sum of “Coulomb-14”, Coulomb short range and Coulomb long range forces, “van der Waals” is the sum of “LJ-14” and van der Waals short and long range terms, and “Nonbonded” is the sum of “Electrostatic” and “van der Waals” terms.

We examine three statistical measures to describe the deviations between the programs: the average differences from the program average over all molecules, the average of the absolute value of the difference from the program average, and the average of the relative absolute value of the difference. The program average is defined as:

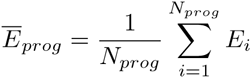

Where the sum is over molecular simulation programs of interest, and *E*_*i*_ is the energy term of interest from the *i*th simulation program. The average difference from the program average is:

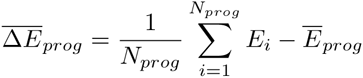

The average of the absolute value of the difference from the program average is defined as:

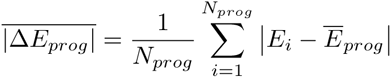

The average of the relative absolute value of the difference is defined as:

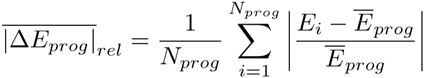

Each statistical measure gives somewhat different information about the trials. 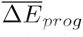 gives information about the manner in which energy component deviates from the program average, while the absolute value of the differences from the average 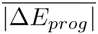 shows the magnitude of the deviation, avoiding any cancellation due to different signs. Because many terms are much smaller than others, for example, the bonded energy terms being two to three orders of magnitude smaller than the electrostatic term, the relative absolute difference 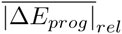 shows the fractional error in each term.

We find that the bonded terms match very well between all programs. Average differences from the program average are below 0.05 kJ/mol in magnitude for all terms for all programs, and usually about an order of magnitude lower, and around or below 0.002 kJ/mol (CHARMM was slightly higher than the others). Average absolute differences in the total bonded term are below 0.004 kJ/mol for all programs except CHARMM. CHARMM’s average absolute value is 0.007 kJ/mol, but is dominated by differences in the dihedral terms; bond and angle terms are as low as other programs. Note that this is not per interaction, but the sum over all interactions.

The average relative absolute quantities are perhaps a more important comparison metric, since they are intensive quantities. Total deviation of the energy will of course become larger as the system becomes larger, so normalizing by the total energy, which will be proportional to system size, will result in a more useful comparison. With this statistic, we see that the bonded terms are accurate to generally about 3 parts in 10^6^, with CHARMM slightly higher at 7 parts in 10^6^. Given that this is approximately the limit of precision one would see in single precision calculations, and given the fact that some programs only output energies to four (AMBER) or five (CHARMM) decimal places, or eight significant digits total (LAMMPS) this seems for all programs to be a reasonable amount of agreement in bonded interactions for most purposes. It demonstrates that the conversion process has successfully copied parameters with the correct functional form for bonded interactions between all of the programs of interest, and the energies are being calculated in a consistent way for these bonded interactions. More generally, it suggests that with no extra fiddling, all programs should in typical cases generate essentially equivalent bonded energies.

We next examine the nonbonded interactions. Coulomb 1-4 and van der Waals 1-4 interactions are a good measure of whether the nonbonded parameters are being copied correctly, as they generally are all calculated with real space interactions and are shorter range than any reasonable cutoff. Therefore, their comparison is not affected by nonbonded simulation parameter choices such as treatment of long-range electrostatics and represents the best test of Lennard-Jones interactions and charges are properly copied from one set of files to another.

We see that, like the bonded interactions, the 1-4 interactions, when separated out from other interactions by the simulation program, are in good agreement. In all available cases, the van der Waals 1-4 interactions have relative absolute differences at least a factor of 2 better than even the bond and angle interactions, at about 1 part in 10^7^. LAMMPS and CHARMM do not calculate 1-4 interactions independently, but some post-processing tricks involving subtracting energies with different input parameters show that the CHARMM van der Waals 1-4 energies have similar accuracy, in particular being generally within output precision of AMBER.

The story from the Coulombic 1-4 interactions is more complicated. Looking at the absolute difference, we see that the difference of GROMACS and DESMOND from the program average is about half of what the difference is from AMBER. In this case, the difference in Coulombic 1-4 interactions between GROMACS and DESMOND is actually less than 10% of what the differences is between AMBER and the other two programs is. Since the program average is the average of the three programs, this means that essentially all the deviation from the program average is because of AMBER’s difference from the other two programs. Since the LJ-14 parameters are shown to be in good agreement by the fact that the LJ-14 energies are in good agreement, the difference must come from some other source.

After further analysis of the data, it became clear that the value of Coulomb’s constant 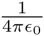, the constant of proportionality *f* in 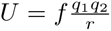 is the main cause of the differences in the Coulomb 1-4 terms. In Table III, we show the value of Coulomb’s constant in a range of different simulation programs compared to the NIST 2014 CODATA value. We list the AMBER number as“AmberTools” because in AMBER, the constant is set by multiplying the charge by 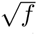 in the `.prmtop` file, rather than set internally by either `sander` or `PMEMD`, the AMBER molecular dynamics engine. Clearly, AmberTools, and to a lesser extent CHARMM, have significant deviation from the best experimental value. It is possible that this difference may primarily come from decisions in the 1970’s to use Bohr’s radius to calculate Coulomb’s constant assuming a value of 0.529167 Å(versus an improved value of 0.592177 Å today), but there appears to be no clear historical answer. Of course, the more important question is, how much does this deviation affect the results?

**TABLE III:**
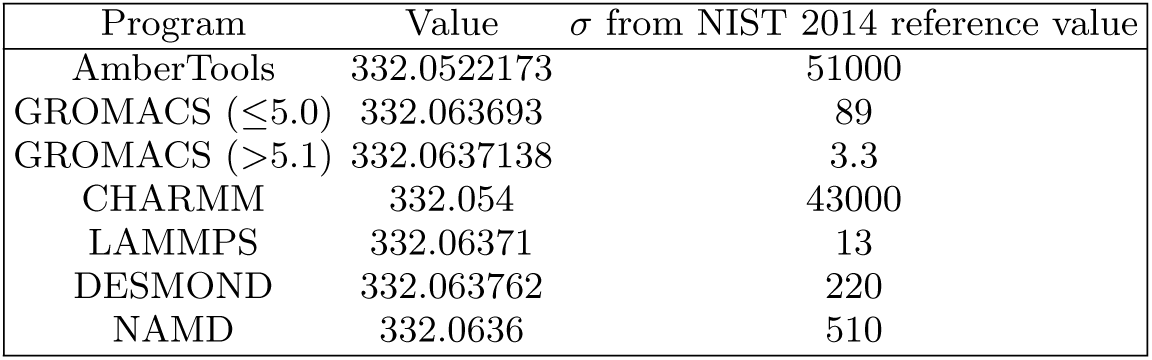
Values of Coulomb’s constant *f* currently used in molecular simulation programs compared to the value of 332.06371302(32) kcal/mol Åe^−2^ or 138.93545753(14) kJ/mol nm e^−2^, calculated from NIST 2014 CODATA. Specific versions of the programs used are described in the text. Two versions of GROMACS are listed because SAMPL5 energies were originally generated with version 5.0.4, but the value has been changed since then. Coulomb’s constant was calculated as *k_e_N_A_e*^2^, where *k*_*e*_ is Coulomb’s constant defined exactly in N m^2^C^−2^ (or J m C^−2^), *N*_*a*_ is Avogadro’s number, and *e* is the elementary charge from NIST 2014 CODATA, and then converted to kcal/mol Åe^−2^. Non-SI units are used in this case because the number represents the value actually coded in the majority of programs. Uncertainties in *f* were calculated using standard error propagation using NIST 2014 CODATA values at the correlation coefficient between *N*_*A*_ and *e* of −0.9985, also from NIST 2014 CODATA. At less than 1000 standard deviations, however, errors due to Coulomb’s constant are no longer the largest source of error.

We tested the effect of changes in Coulomb’s constant on the energy. To add a more rigorous control, we looked at the RMS difference in energy between AMBER energies and GROMACS energies evaluated with the Coulomb constant from version 5.0.4, and then with GROMACS recompiled with the AmberTools Coulomb’s constant. Results are shown in Table IV, where we have used RMSD over all molecules to compare these programs. We see that matching Coulomb’s constant removes 98.8% of the variation in the Coulomb 1-4 term between the two programs, 69.5% of the total electrostatic energy difference, and 74.2% of the total potential energy difference between the two programs, strongly indicating that the lack of agreement of AMBER with the other programs is almost entirely a result of mismatched Coulomb’s constant.

**TABLE IV:** RMSD in kJ/mol of different energy components in GROMACS 5.0.4 from AMBER energies as the GROMACS Coulomb’s constant is varied. Averages are calculated over all 22 SAMPL5 host-guest systems.

|  | Coulomb-14 | Electrostatic | Total Potential |
| --- | --- | --- | --- |
| GROMACS original constant | 0.00522 | 0.789 | 0.872 |
| AmberTools constant | 0.000064 | 0.241 | 0.225 |
| Percent difference explained | 98.8% | 69.5% | 74.2% |

The longer range nonbonded interactions are significantly harder to get in good agreement between programs. Validating the van der Waals and Coulombic 1-4 interactions demonstrates that the Lennard-Jones parameters and Coulombic charges are correctly created in the other file formats. In that sense, validating the conversion of file formats can be done without comparing the long-range interactions. However, if we are interested in comparing results of molecular dynamics programs in realistic situations, we will need to compare the entire potential energy, including these terms.

One complication is that different programs both calculate and print out the different components of nonbonded interactions differently, such as the direct space energy, the Fourier space energy, the Ewald self term, and so forth. Thus, it is often difficult to examine anything except the total Lennard-Jones or total electrostatic energy. This makes it hard to determine exactly the source of any discrepancy between programs, and motivates our attempt to find ‘ideal’ simulation parameters to best make this comparison.

Discrepancies in the nonbonded energy terms are always much larger in magnitude than discrepancies in the bonded interactions for any liquid phase simulation, since there are many more intermolecular interactions than intramolecular interactions. It is therefore instructive to look first at the average relative absolute deviations from the program average (Figure 1). For GROMACS, AMBER, LAMMPS, and CHARMM, the fractional difference in the van der Waals energy is approximately 1 *×* 10^−5^, which is 2-5 times larger than the difference in the bonded energy (DESMOND does not separate the long-range energy out into components). GROMACS and LAMMPS are generally closer to each other, usually within one part in 10^6^, and CHARMM and AMBER are clustered together, though not as closely as GROMACS and LAMMPS. Because of the close match of Lennard-Jones 1-4 parameters, deviations in the total Lennard-Jones energy are likely due to differences in the calculation of long range interactions.

**FIG. 1:**
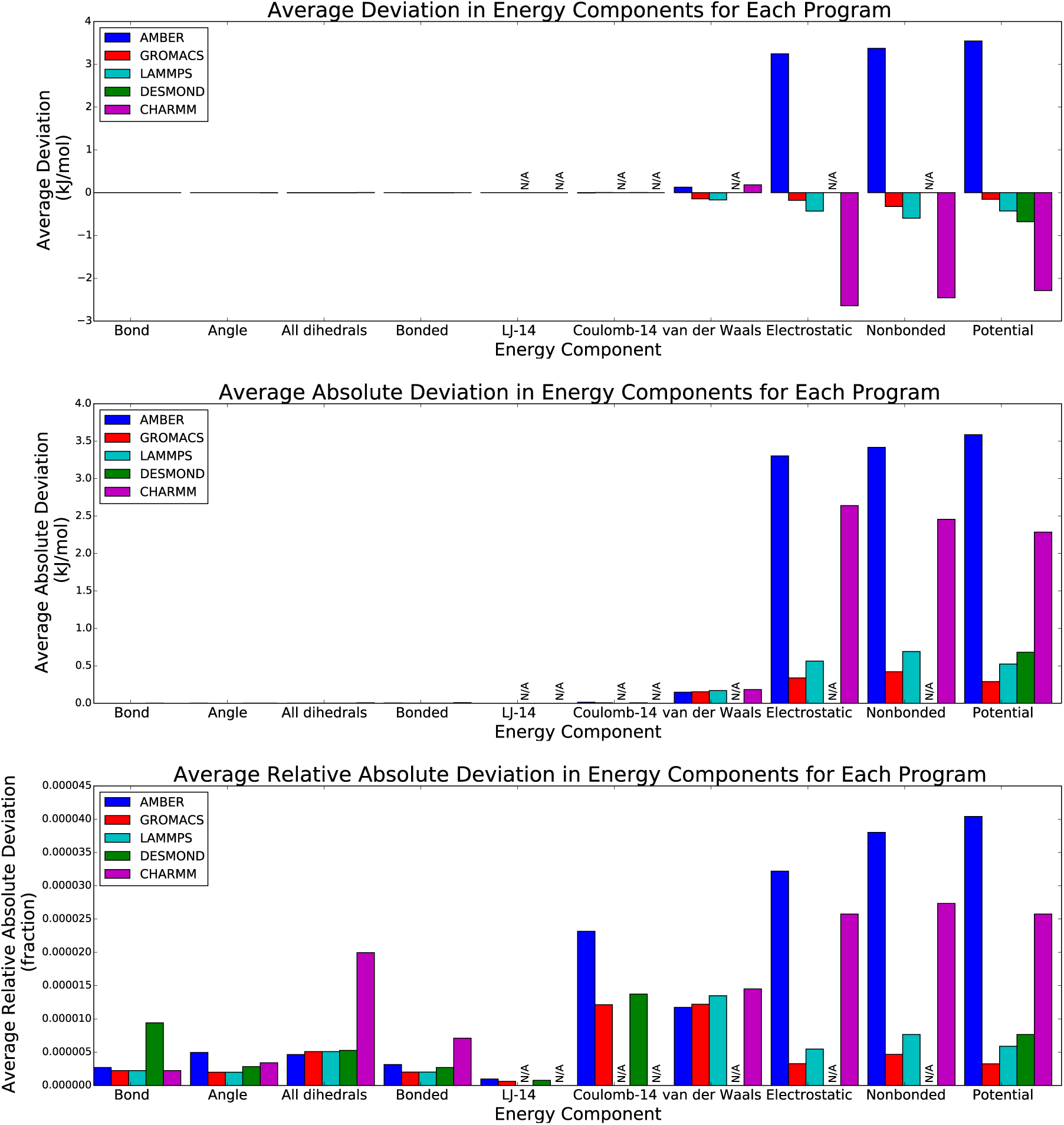
Electrostatic energy interactions dominate the deviations between the programs both in absolute and relative error, though many components have measurable relative error. We compare the variation of 10 different energy terms between five different simulation programs (AMBER, GROMACS, LAMMPS, DESMOND, and CHARMM) for the ‘ideal’ choice of cutoff parameters. For each term, we plot the deviation of each program from the average over all programs (the program average), to avoid choosing a single arbitrary reference program. All statistical measures are averaged over the 22 SAMPL5 host-guest systems. We plot the average deviation (top), the absolute average deviation (middle), and relative absolute average deviation (bottom). ‘N/A’ is listed when the energy term cannot be extracted from the simulation output for that program.

Differences from the program average in the total electrostatic energy are smaller than errors in the total van der Waals energy for GROMACS and LAMMPS, but are significantly larger for AMBER. As seen above, a large portion of the electrostatic deviation of AMBER from those two programs is because of the inconsistent choice of Coulomb’s constant. Although CHARMM also has a relatively inaccurate Coulomb’s constant, the differences in the CHARMM potential energy are of opposite sign from AMBER, indicating a difference in the way the electrostatic energy is calculated relative to the other programs. For AMBER, about 70% of this deviation is due to Coulomb’s constant choice, leaving significantly less of the deviation from other programs due to other choices in nonbonded simulation parameters. Correcting CHARMM’s Coulomb’s constant would actually make the energy further from the other three programs. However, it is important to notice that the fractional difference is still on the order of 2.5 × 10^−5^, likely too small to matter for most quantities of interest over long simulations.

The deviations from the program average of the total potential energy are dominated by the differences in the nonbonded terms, since those are so much larger than differences in the bonded energy. For CHARMM and AMBER, differences in the electrostatic nonbonded energy dominates. DESMOND total potential energy differences from the program average are almost the same as GROMACS and LAMMPS, indicating that since the bonded interactions match the two programs well, the nonbonded also must agree relatively well.

The energies generated using the ‘default’ parameters defined above are shown in Figure 2. The bonded interactions are essentially identical, as there are no significant differences between the simulation parameters used for the ‘ideal’ and ‘default’ energy evaluations. AMBER, GROMACS AND LAMMPS nonbonded deviations from the program average are approximately twice as large with the default states, while DESMOND nonbonded deviations are approximately five times larger. The patterns are similar to what is seen with the ‘ideal’ parameters, but the larger magnitudes obscure the sources of these deviations in energy, requiring the creation of the ‘ideal’ parameter set. In most cases of simulating physical observables, these deviations would still likely be accurate enough for most thermodynamic calculations, with relative average absolute difference of 0.005% from each other, likely much smaller than the errors due to statistical variation in the calculation itself.

**FIG. 2:**
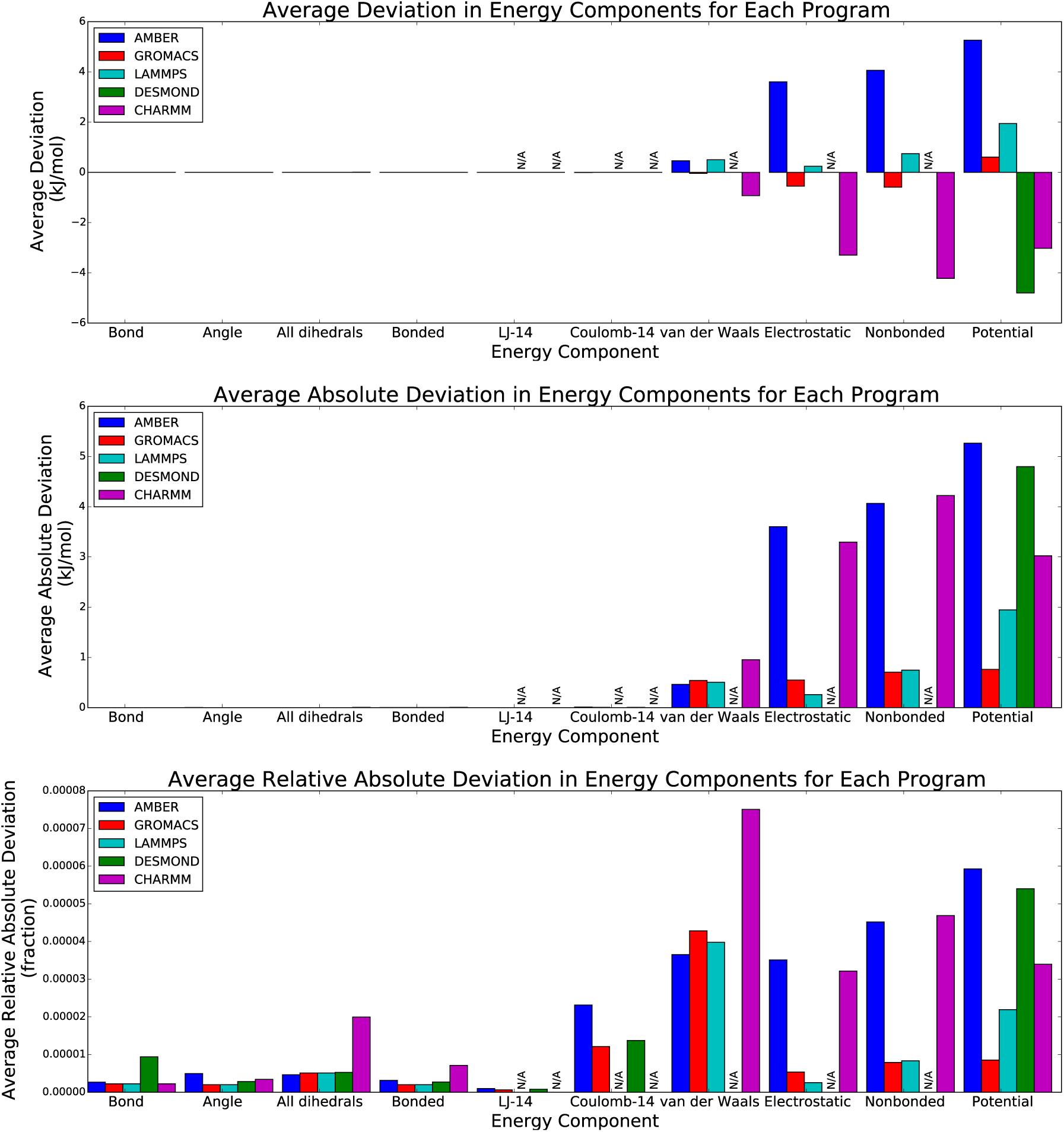
We compare the variation of 10 different energy terms between five different simulation programs (AMBER, GROMACS, LAMMPS, DESMOND, and CHARMM) for the ‘default’ choice of cutoff parameters (described in Table II). As above, for each term, we plot the deviation of each program from the average of all programs, to avoid choosing a single arbitrary reference program. All statistical measures are averaged over 22 molecules. We plot the average deviation (top), the absolute average deviation (middle), and relative absolute average deviation (bottom). Nonbonded potential parameter deviations are approximately a factor of 2–5 larger than using the ‘ideal’ parameters. ‘N/A’ is listed when the energy term cannot be extracted from the simulation output for that program.

We also are interested in the deviations of energies as a function of the number of digits used in the coordinate output file in order to determine what is required to convert between different simulation formats with high fidelity. The results of the comparison between AMBER (full precision input) and GROMACS (reduced precision output) are shown in Fig. 3. We choose only to show GROMACS, as the other programs show similar behavior. We plot the − log_10_ of the average relative absolute difference between the GROMACS energy and the AMBER energy as a function of the number of decimal places in the output GROMACS coordinates from 8 digits after the decimal place (measured in nanometers) down to 4 after the decimal place, what one would obtain from a file downloaded from the PDB.

**FIG. 3:**
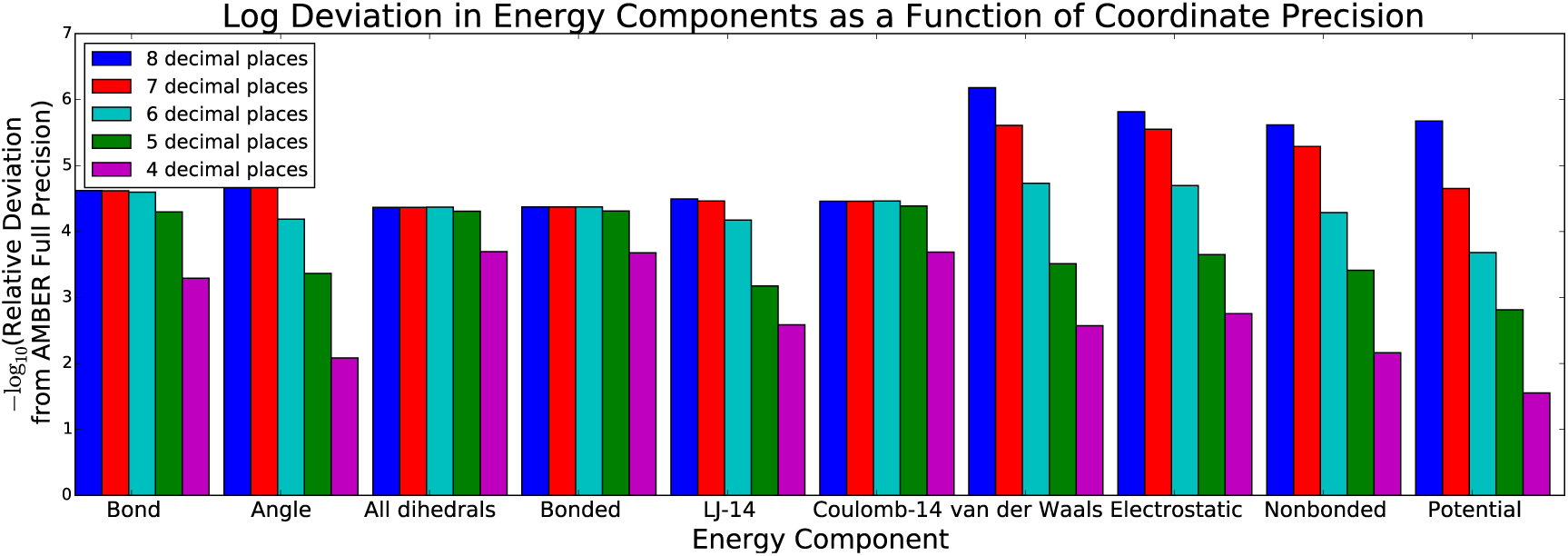
Matches in energy between converted files become rapidly worse as the number of digits of precision in the converted coordinate files decreases. We plot − log_10_ of the average relative absolute value in each energy term between AMBER and GROMACS, with fixed input coordinate precision, and variable output precision with the number of decimal places in the coordinates in nanometers, varying from 8 down to 4, the precision of a standard PDB file.

We see that the total energy loses approximately 0.75-1 digits of relative precision in the energy for each digit of precision of coordinate removed. Since the van der Waals and electrostatic nonbonded energies contribute the majority of the potential energy, their loss of precision mirrors the overall loss of precision. Interestingly, the bonded and both 1-4 terms are less sensitive to changes in coordinate precision, not losing much precision until getting down to 5 or fewer digits after the decimal point. Bond and angle terms individually lose precision, but the dihedral energy dominates the total bonded energy. Both Lennard-Jones and electrostatic nonbonded energies fall off similarly in precision, so large errors created by rounding in the repulsive *r*^−12^ interactions do not seem to be larger than changes in the Coulombic *r*^−1^ terms as the coordinates become more approximate. Overall, losing just a few digits of precision completely washes out any other source of error observed in this study, demonstrating the importance of matching the coordinates to high precision in order to validate the rest of the conversion.

We are also interested in how much changing the numerical precision of the energy calculation affects energy comparisons. We focus on the comparison between single and double precision GROMACS, as it is specifically designed to be compiled in either single and double precision, though the single precision is the default version most simulations are run with. In Table V, we compare the RMS differences averaged over all 22 compounds between the two GROMACS binaries, AMBER and single precision GROMACS, and AMBER and double precision GROMACS. We use RMS differences instead of average deviation statistical measures presented above as the effects of precision are most likely to affect the variation of the results, rather than the average deviations. This comparison allows us to see both the magnitude of the difference due to changes in binary precision, and how much this differences affects the comparison to, for example, AMBER.

**TABLE V:** Differences between double and single GROMACS energy evaluations are of similar magnitude to the differences between AMBER and GROMACS, but are dominated by differences in the long-range electrostatics. All energies in kJ/mol.

| E term | RMS($E_{single} - E_{double}$) | RMS($E_{amber} - E_{single}$) | RMS($E_{amber} - E_{double}$) |
| --- | --- | --- | --- |
| Bond | 0.000066 | 0.000068 | 0.000008 |
| Angle | 0.000044 | 0.000043 | 0.000007 |
| All dihedrals | 0.000015 | 0.000031 | 0.000018 |
| Bonded | 0.000081 | 0.000086 | 0.000011 |
| LJ-14 | 0.000013 | 0.000020 | 0.000025 |
| Coulomb-14 | 0.000007 | 0.001250 | 0.001251 |
| van der Waals | 0.001894 | 0.021756 | 0.023116 |
| Electrostatic | 0.218874 | 0.403839 | 0.189265 |
| Nonbonded | 0.217209 | 0.422827 | 0.209207 |
| Potential | 0.217134 | 0.422781 | 0.209214 |

The differences between single and double precision shown in Table V for bonded terms are 2-8 times larger than the difference between AMBER and double precision GROMACS, and are usually the dominant contribution to the difference between AMBER and single precision GROMACS. Differences between the precisions for LJ-14 are about equal to differences between either precision and AMBER. Differences between precisions for Coulomb-14 terms are as low as the difference between LJ-14, which is, of course, much less than the GROMACS to AMBER difference, because of the previously described difference in Coulomb constant. The differences in precision for the total van der Waals energy is an order of magnitude lower than the difference between AMBER and either GROMACS precision caused by differences in calculating the long-range nonbonded terms. However, the single to double precision change results in significant difference in the overall electrostatic term, as large as the magnitude between GROMACS and AMBER. Because the total van der Waals energy changes relatively little between precisions, it is likely that the short-range electrostatics (which are functions only of the distance) are also relatively accurate, and it is the Ewald summation part that changes upon changes in binary precision.

We are also interested in how much of the differences between programs vary with the configurations of each molecule. For example, if we were to take different configurations of the same molecule, would we get similar deviations from the program average for all of the molecules? We address this question by taking the 12 octa-acid hosts, and generating 20 configurations as described in the Methods section using NVT molecular dynamics. The average over these 20 configurations is a rough approximation to the ensemble average energy of the system.

We then compare the RMSD from the program average *σ*_*config*_, averaged over all 12 *×* 20 = 240 configurations, and the RMSD of the average energy of all configurations of the same molecule from the program average over the 12 host-guest systems *σ*_*molecule*_. If the variation from program to program is independent of configuration, and only dependent on the differences between molecules, then we would expect that the two different RMSDs would be roughly equal, meaning there is low conformation dependent variation. If instead the variation is independent of the specific molecule, and depends on interactions from random atoms, independent of the molecule, then the RMSD from the configurationally averaged deviations from program averages,*σ*_*molecule*_, would be significantly smaller (approximately 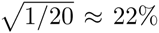 of the value *σ*_*conformation*_ value). The extent to which *σ*_*molecule*_ is smaller than *σ*_*config*_ shows how much of the variation is inherent to the molecules, and how much is only dependent on the configurations.

We can quantify this difference in the source of variation by calculating the fraction of the total variation due to conformational variability, calculated as 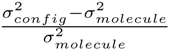, for each energy term. If this quantity is low, then variation is mostly due to differences between molecules, not configurations. If it is near one, then variation between programs is mostly due to changes in conformation.

We can observe the results in Figure 4. At one extreme are the bond energies, which have only about 8% of the total variation between programs due to configurational variation, near the minimum of 1/20 *≈* 5%. Most of the differences are due to differences between the molecular bond terms, but not the specific conformation. Similarly, the variations in van der Waals 1-4 interactions are mostly due to differences between molecules.

**FIG. 4:**
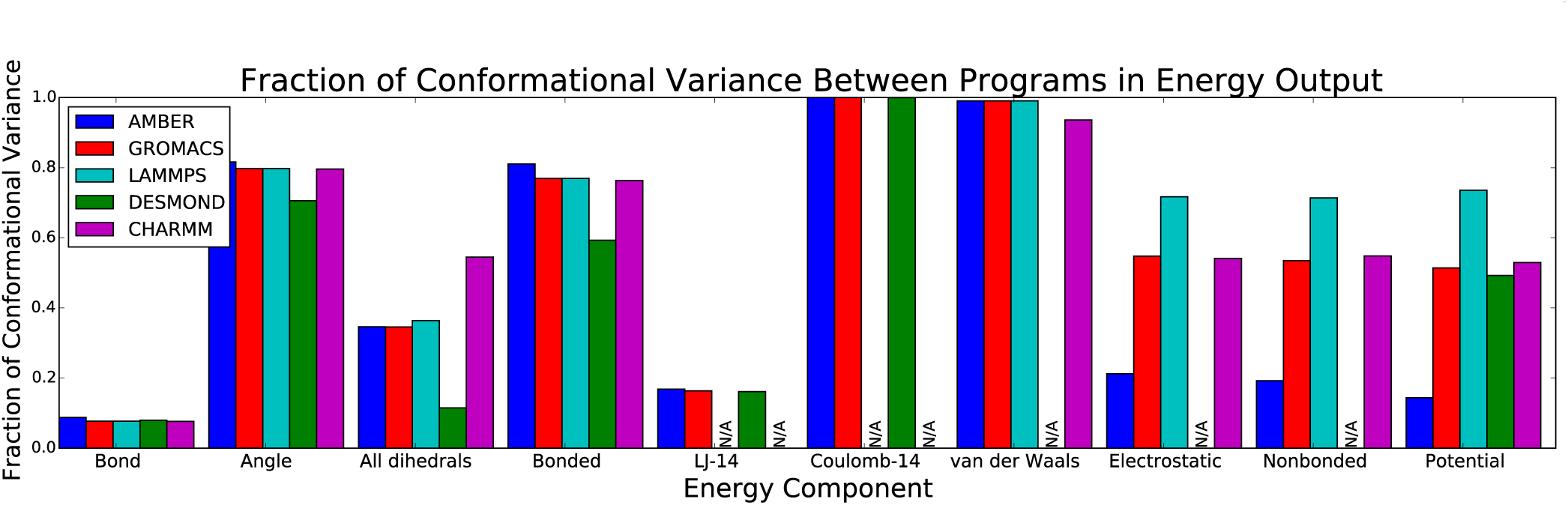
Fraction of variation from the program average due to conformational variability instead of molecular variability, calculated as 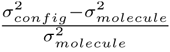, where *σ*_*config*_ and *σ*_*molecule*_ are standard deviations from the program energy over all configurations and over molecule averages, respectively.

At the other extreme are the Coulomb 1-4 terms, where almost 100% of the variation is due to conformational variation: after averaging the differences over molecules, there is very little variation left due to conformational changes. It is not entirely clear why Coulomb 1-4 and van der Waals 1-4 are so different in patterns, given that they are both primarily determined by the locations of the same set of 1-4 atoms. Similarly, almost 100% of the variability in the total van der Waals energy is due to conformational variability. Variation in angle energies from the program average is again relatively dependent on configuration (around 80%).

Total electrostatic energy variation is one of the few energy components where the fraction of conformational variation depends significantly on the programs. For AMBER, it is not very dependent on configuration; for other programs, it is much more dependent on configuration. This is likely because of differences in Coulomb’s constant and in the treatment of long-range electrostatic energies; Coulomb’s constant changes can make such a difference because the long-range energies are much less dependent on individual molecular distances, instead being dependent on the average distribution of charge within the system, which does not change significantly for a host-guest system over time, and which will scale with the changes in Coulomb’ constant. On the other hand, the Coulomb 1-4 interactions are dominated by the variability of which atoms are closest to each other at any given time, which is larger than the Coulomb constant differences.

The analysis in the variation of the total potential energy illustrates again that the dominant reasons that the molecular simulations differ are the evaluation of long-range interactions, especially the electrostatics, and the choice of the Coulomb constant. We find that the total variation of the potential energy, like total potential energy itself, depends almost entirely on the nonbonded terms. Since the van der Waals variation between programs is almost entirely conformation dependent, with very little deviation in programs between the ensemble average estimate for each molecule, the conformational dependence of the total energy is essentially determined by the conformational dependence of the electrostatic energy. The differences in bonded terms are essentially negligible.

## DISCUSSION

### Issues in file conversion

Creating true all-to-all functionality, from any of a set of molecular simulation programs to any other, is particularly difficult. There are a number of one-to-one conversion utilities and scripts: for example, ACPYPE [12], CHAMBER [13], and `amber2lmp`. InterMol and ParmEd have been explicitly developed as all-to-all converters, though not all functionality yet exists. A truly all-to-all converter would substantially simplify the current painstaking process of molecular simulation conversion.

There are a number of differences between programs not immediately obvious that nonetheless need to be carefully matched for the same system to be represented in both programs. For example, GROMACS builds lists of 1-4 interactions based on the bond topology: if three bonds connect two atoms, they are 1-4 interactions. However, AMBER uses the presence of dihedrals to define 1-4 interactions. A dihedral with zero energy in GROMACS is essentially redundant and can be eliminated without affecting the energy, but is used to define 1-4 interactions in AMBER. Additionally, the same interaction in different programs may have different names and functional formats. This can either be handled by hard-coded conversions (ParmEd) or converting all interactions through a canonical form (InterMol). The first has the advantage of readability, whereas the second is more general. It is not yet clear what the optimal strategy would be.

One of the most challenging problems in conversion of molecular simulation files is handling the different units in each simulation engine. Both InterMol and ParmEd address this problem by automatically converting units between simulation input files, removing the need for manual unit conversion. This is handled by creating data types that carry units with them, making conversion much simpler. Without such unit-carrying data types, doing anything other than one-to-one conversion becomes significantly harder.

### Issues in matching energies

It is difficult to say what the “right” energy is for a given force field, as there are several different reasonable choices for implementation of long-range interactions. For a sufficiently large box, one could simply extend out the cutoffs, treating an increasingly larger amount of the system using straightforward direct space interactions. However, these systems, at 4.0 nm across, are not large enough to take the direct-space electrostatic treatment out far enough for the different programs to completely converge together. This remains a key weakness of the study, making it difficult to decide on a single reference energy to compare the simulations. This resulted in our choice to examine deviations from the program average, rather than attempt to determine the correct energy.

One could state that the differences described between the programs (what switching scheme is used, what cutoff is used, Coulomb’s constant), since these properties influence the parameterization of the force field, should actually be considered as part of the force field to reproduce simulations. However, explicitly stating all possible variants of simulations that might affect the energy would require significant additional functionality added to each different molecular simulation engine. Instead, it is preferable to implement methods such as periodic summation of Lennard-Jones or long range corrections [18, 25] that result in simulation parameters such as the cutoff and switching scheme not affecting the total energy or force in a statistically significant way. This way, the developers of each simulation engine can choose the parameter-independent algorithms that best suited their needs.

In order to aid further analysis of the energies, all energy output and summarized analysis presented here in this paper is provided in the ‘analysis.tgz’ directory in the supporting information, as described in the ‘README.txt’ supporting information file.

## CONCLUSIONS

The results from this study confirm the simulation input files for SAMPL5 are properly converted, based on the very strong agreement, essentially within single precision rounding, for all bonded terms, Lennard-Jones 1-4 terms and (with one caveat outlined below) Coulomb 1-4 interactions. Only the long-range interactions deviated by moderate amounts between programs, especially the electrostatic Fourier-space interactions.

The results also strongly suggest that all molecular simulation programs should choose a sufficiently consistent value for Coulomb’s constant. A value that deviates from experiment by 0.01 out of 332.06371 kcal/mol Åe^−2^ (one part in 10^5^) is simply not accurate enough to allow simulations to agree. We find that that once the deviation is below 0.0001, this error contributes less than other sources of error, so a value of 332.06371*±*0.00005 should reduce this sort of discrepancy to a negligible error. This level of agreement is present in all current programs with the exception of CHARMM and AMBER, with GROMACS, at the edge of that range, recently improving the precision by a factor of 4 in the 5.1 release.

Other than the difference in Coulomb’s law, most programs agree quite well, probably enough for most practical purposes as most thermodynamic quantities cannot be measured experimentally that accurately. Even those differences are unlikely to affect most thermodynamics computations to within the statistical error generally accepted in simulation studies. The energy calculations for all molecular dynamics engines for this molecular set agree to a better than 0.1% relative absolute energy for all energy components, and in most cases an order of magnitude better, when reasonable choices are made for different cutoff parameters. However, to get good agreement, we must use carefully matched cutoff treatments, as different ways to shift or switch the cutoffs can introduce significant differences between the energies of the same system evaluated with different simulation engines.

We believe that this study represents the largest automated comparison between different programs that has been performed so far, a process only possible because of the automated conversion. The pipeline described in this program, properly scaled up for diverse ranges of systems, could be useful for a range of comparison studies that require the precise comparison across multiple simulation engines, or in modeling pipelines that require one simulation program for part of the pipeline, and another simulation engine for a different part.

## ACKNOWLEDGMENTS

The authors would like to thank Frank Pickard (NIH) for sample CHARMM inputs and discussion about evaluation of CHARMM energies, Justin Lemkul (U. Maryland) for advice on CHARMM functional form, and Chris Lee (U. Va., UCSD), Alex Yang (U. Va.), Michael Zhu (U. Va.), Hari Devanathan (U. Va.), and Jacob Rosenthal (U. Va.) for initial work on InterMol. DLM thanks the NSF (CHE 1352608) and the NIH (1R01GM108889) for financial support. MKG thanks the NIH (R01GM061300 and U01GM111528) for financial support. DAC thanks the NIH (R01GM045811) for financial support.

